# Insights into the mitochondrial transcriptome landscapes of two Brassicales plant species, *Arabidopsis thaliana* (var. Col-0) and *Brassica oleracea* (var. botrytis)

**DOI:** 10.1101/2020.10.22.346726

**Authors:** Corinne Best, Laure Sultan, Omer Murik, Oren Ostersetzer-Biran

**Affiliations:** Department of Plant and Environmental Sciences, The Alexander Silberman Institute of Life Sciences, The Hebrew University of Jerusalem, Givat-Ram, Jerusalem 91904, Israel; Medical Genetics Institute, Shaare Zedek Medical Center, Jerusalem, Israel

**Keywords:** mtRNA, transcriptome, group II intron, splicing, Mitochondria, Brassicales

## Abstract

Mitochondria play key roles in cellular energy metabolism within eukaryotic cells. As relics of endosymbiotic bacteria, most (but not all) mitochondria contain their own genome (mitogenome, mtDNA), as well as intrinsic biosynthetic machinery for making organelle RNAs and proteins. The expression of the mtDNA requires regulated metabolism of its transcriptome by nuclear-encoded factors. Post-transcriptional mtRNA modifications play a central role in the expression of the plant mitogenomes, and hence in organellar biogenesis and plant physiology. Despite extensive investigations, a full map of angiosperm mitochondrial transcriptomes, a prerequisite for the elucidation of the basic RNA biology of mitochondria, has not been reported yet. Using RNA-seq data, RT-PCR and bioinformatics, we sought to explore the gene expression profiles of land plant mitochondria. Here, we present the mitochondrial transcriptomic maps of two key *Brassicaceae* species, *Arabidopsis thaliana* (var Col-0) and cauliflower (*Brassica oleracea* var. botrytis). The revised transcriptome landscapes of Arabidopsis and cauliflower mitogenomes provide with more detail into mtRNA biology and processing in angiosperm mitochondria, and we expect that they would serve as a valuable resource for the plant organellar community.

**Accession numbers:** Sequences are available at the Sequence Read Archive (accession no. PRJNA472433), for both *Arabidopsis thaliana* var. Col-0 mtRNA (SRA no. SRX4110179) and *Brassica oleracea* var. botrytis mtRNA (SRA no. SRX4110177).

## Introduction

Mitochondria house the oxidative phosphorylation (OXPHOS) machinery and various other essential metabolic pathways in the plant cell (Millar *et al*. 2011, Schertl and Braun 2014). As descendants from an ancestral proteobacterial symbiont, the vast majority of the mitochondria contain their own genetic system (mitogenome, mtDNA), which encodes for rRNAs and some of the organellar tRNAs and proteins (Bonen 2018a, Gualberto and Newton 2017). Plants are able to coordinate their energy demands during particular growth and developmental stages by means of nucleocytoplasmic signaling (*i*.*e*. nuclear to mitochondria or plastids). The metabolic functions, biogenesis and maintenance of the mitochondria are controlled by complex networks of genetic interactions between the host genome and the organelles, activities which may relate to plant terrestrialization, about half a billion years ago (Best *et al*. 2020). For instance, the mitochondrial ribosomes and the energy transduction machineries are assemblies of both nuclear and organellar encoded subunits, were the correct stoichiometry in the accumulation of the different subunits composing the organellar complexes is necessary for their biogenesis and functions. These processes necessitate complex mechanisms for regulating the coordination of the expression and accumulation of the different subunits encoded by two physically remote genomes, involving both anterograde and retrograde signaling cascades (Fuchs *et al*. 2020, Kleine and Leister 2016, Woodson and Chory 2008). However, the identity of the messenger molecules involved in these pathways is more elusive.

The use of high throughput sequencing technologies has revolutionized the field of organelle genomics and genetics. Among the main findings are recent indications of the extreme complexity and variability of land plant mitogenome structures, gene expression and RNA metabolism patterns (Colas des Francs-Small and Small 2014, Hammani and Giege 2014, Zmudjak and Ostersetzer-Biran 2017). Although all mitochondria are assumed to share a common ancestor (Gray *et al*. 2001), these have diverged considerably in different clades. For example, the protist *Reclinomonas Americana* seem to contains the most bacteria-like organellar genome known to date (Burger *et al*. 2013), whereas in the dinoflagellate *Amoebophrya ceratii*, the entire mitogenome was lost and the essential component of the respiratory apparatus were all translocated into the nuclear genome (John *et al*. 2019).

In animals, the mitogenomes are typically small and conserved circular-mapping molecules of *ca*. 16 kbp, encoding 37 tightly packed genes that are organized in two main transcription initiation sites, known as the light- and heavy-strand promoter sites (*i*.*e*. LSP and HSP, respectively) (Clayton 1991). The mtDNAs in land plants seem notably larger and more variable in size, sequence and structure (Bonen 2018a, Gualberto and Newton 2017). These may also contain linear or circular extrachromosomal DNAs, and seem to undergo frequent recombination events that lead to rapid changes in genome configuration, and also result in the formation of many novel open reading frames (ORFs), many of which have no assigned functions (Bonen 2018a, Gualberto and Newton 2017). Some plants have experienced dramatic expansion in mitogenome size, resulting in the largest known mitochondrial genomes (*i*.*e*. ∼11.3 Mbp) (Sloan *et al*. 2012). Others, as the mitochondrial genome in mistletoe (*Viscum album*) undergone massive gene loss (Maclean *et al*. 2018, Petersen *et al*. 2015, Senkler *et al*. 2018). Despite the large variation in mitogenome size and gene organization, the number of mitochondrial genes is relatively conserved in the land-plant kingdom, with about 60 known genes found in different terrestrial plant species (Bonen 2018b, Grewe *et al*. 2014, Gualberto and Newton 2017, Guo *et al*. 2016, Mower *et al*. 2012, Park *et al*. 2015, Sloan, *et al*. 2012). These include tRNAs, rRNAs, ribosomal proteins, various subunits of the respiratory complexes I (NADH dehydrogenase), III (cytochrome c reductase or bc1), and IV (cytochrome c oxidase), subunits of the ATP-synthase (also denoted as CV), cytochrome *c* biogenesis (CCM) factors, and at least one component of the twin-arginine protein translocation complex.

The transcription of the mtDNA in animals is initiated from a small noncoding region, known as the D-loop site (Clayton 1982). The mitochondrial genes are generally transcribed as two large precursor polycistronic transcripts, which are subsequently cleaved to generate individual mRNAs, tRNAs and rRNAs (Mercer *et al*. 2011). The expression of the mitogenome in angiosperms is notably more complex. Different from their counterparts in Animalia, the mitochondrial genes in land plants are arranged in numerous polycistronic units, which are physically separated by relatively large and non-conserved intergenic regions (Zmudjak and Ostersetzer-Biran 2017). These are postulated to harbor important regulatory regions, as promoters, enhancer or repressor DNA sequences, which can activate or downregulate the expression of organellar genes in a tissue, developmental or environmental dependent manner. Post-transcriptional RNA processing plays a pivotal role in plant’s mtDNA expression (Best, *et al*. 2020, Bonen 2018a, Colas des Francs-Small and Small 2014, Hammani and Giege 2014, Zmudjak and Ostersetzer-Biran 2017). These involve the maturation of 5’ and 3’ termini, numerous RNA editing events (typically C-to-U exchanges) in most of the mitochondrial transcripts (Small *et al*. 2020) and the removal of intron sequences that reside within many essential genes (Brown *et al*. 2014, Schmitz-Linneweber *et al*. 2015). These RNA processing steps are essential for the organellar RNAs to carry out their functions in protein synthesis, and may therefore serve as key control points in plant mitochondria gene expression.

The processes that lead to the establishment of functional mRNAs in the mitochondria are accomplished largely by nuclear-encoded cofactors, which may also provide a means to link organellar functions with environmental and developmental signals. The nuclear genomes of angiosperms encode numerous RNA binding proteins that are postulated to reside within the mitochondria (Colas des Francs-Small and Small 2014, Zmudjak and Ostersetzer-Biran 2017). These enzymes comprise a staggering proportion of the total proteome of angiosperm mitochondria, representing about 15% of the identified protein species (Fuchs, *et al*. 2020), further signifying the importance of RNA metabolism for mitochondrial biogenesis and plant physiology (Best, *et al*. 2020). In spite of these data, a detailed map of the mitochondrial RNA landscapes of angiosperm’s mitochondria has not been provided yet. Here we used RNA sequencing and gene expression resources to investigate, at nucleotide resolution, the transcriptomic landscapes of mtRNA metabolism profiles of two key Brassicales species, *Arabidopsis thaliana* (var Col-0) and cauliflower (*i*.*e. Brassica oleracea* var. botrytis), with an aim to establish their relevance to mitochondria biogenesis and plant physiology.

## Material and Methods

### Plant material and growth conditions

Plants growth and analyses generally followed the procedures described in (Sultan *et al*. 2016b). *Arabidopsis thaliana* ecotype Columbia (Col-0) seeds were obtained from the ABRC center, at Ohio State University (Columbus, OH). The seeds were sown on MS-agar plates, incubated in the dark for at least 4 days at 4°C, and then transferred to controlled temperature and light conditions in the growth chambers [*i*.*e*. 22°C under short day conditions (8-hour light, 150 µE·m^-2^·s^-1^ and 16-hour dark)]. Cauliflower (*Brassica oleracea* var.) inflorescences were purchased fresh at local markets and used immediately.

### Mitochondria extraction and analysis

Isolation of mitochondria from 3-week-old Arabidopsis seedlings grown on MS-plates was performed according to the method described in (Keren *et al*. 2012). Preparation of mitochondria from cauliflower inflorescence followed the protocol described in detail in (Grewe, *et al*. 2014, Neuwirt *et al*. 2005, Sultan *et al*. 2016a). In brief, about 1.0 kg cauliflower inflorescences were cut off from the stems, and ground with ice-cold extraction buffer [0.45 M mannitol, 45 mM Na-pyrophosphate, 3 mM EDTA, 1.2% PVP25 (w/v), 0.45% BSA (w/v), 4.5 mM cysteine, 7.5 mM glycine, and 3 mM β-mercaptoethanol; pH 7.5]. Mitochondria were recovered from the extract by differential centrifugations and purification on Percoll gradients, aliquoted and stored frozen at -80^°C^.

### RNA extraction and analysis

RNA extraction from highly-purified mitochondria was performed essentially as described previously (Cohen *et al*. 2014, Keren *et al*. 2011, Zmudjak *et al*. 2017). In brief, RNA was prepared following standard TRI Reagent^®^ protocols (Thermo Fisher Scientific, Cat. no. 15596026). The RNA was treated with RNase-free DNase (Ambion, Cat no. AM2222) prior to its use in the assays. Reverse transcription was carried out with Superscript III reverse transcriptase (Thermo Fisher Scientific, Cat no. 18080093), using 1 µg of total Arabidopsis or cauliflower mtRNA and 250 ng of a random hexanucleotide mixture (Promega, Cat no. C1181) and incubated for 50 min at 50°C. Reactions were stopped by 15 min incubation at 70°C. Reverse transcription PCR (RT-PCR) was performed essentially as described previously (Sultan, *et al*. 2016b, Zmudjak, *et al*. 2017).

### Transcriptome mapping by high-throughput (RNA-seq) analysis

Total mtRNA was extracted from mitochondria preparations, obtained from Arabidopsis and cauliflower inflorescences. Sequencing was carried out by Illumina Genome Analyzer (Center for Genomic Technologies, The Hebrew University of Jerusalem, Israel). Reads were filtered for low quality and primers or adapters contaminations using trimomatic (ver. 0.36) (Bolger *et al*. 2014). Filtered reads were mapped to the Arabidopsis (BK010421) (Sloan *et al*. 2018) and cauliflower (KJ820683) (Grewe, *et al*. 2014) mitochondrial genomes, using Bowtie2 (version 2.2.9) (Langmead and Salzberg 2012). Sequences are available at the Sequence Read Archive (accession no. PRJNA472433), for both *Arabidopsis thaliana* var. Col-0 mtRNA (SRA no. SRX4110179) and *Brassica oleracea* var. botrytis mtRNA (SRA no. SRX4110177).

## Results

### Analysis of the mitochondrial transcriptome landscapes of Arabidopsis and cauliflower

The regulation of mtDNA expression is critical for controlling the OXPHOS capacity in response to physiological demands and environmental signals (Zmudjak and Ostersetzer-Biran 2017). Analyses of the RNA landscapes in plant mitochondria provided with important insights into the complexity of their RNA metabolism and indicated that the regulation of mtRNA processing plays a central role in angiosperm’s mitochondria gene expression (reviewed by e.g. (Zmudjak and Ostersetzer-Biran 2017). Although the importance of RNA processing events to plant mitochondria gene expression and respiratory-mediated functions (and hence for plant physiology), a detailed map of the transcriptome landscapes of angiosperm’s mitochondria has not been provided yet.

*Brassicaceae* (also known as Cruciferae, or mustard family) comprise numerous important crop species, as well as the preeminent model plant *Arabidopsis thaliana*. This study focuses on two key crucifer species, *Arabidopsis thaliana* (var. Col-0) and cauliflower (*Brassica oleracea* var. botrytis). While Arabidopsis serves as a prime model system for plant molecular genetics, cauliflower is employed for biochemical analysis of plant mitochondrial RNA metabolism (Grewe, *et al*. 2014, Neuwirt, *et al*. 2005, Sultan, *et al*. 2016a). To generate the transcriptome maps of Arabidopsis and cauliflower mitochondria, we used total mtRNA extracted from highly enriched mitochondria preparations obtained from Arabidopsis seedlings and cauliflower inflorescence (Murik *et al*. 2019). The mtRNA from each plant species was subjected to Illumina sequencing (see Materials and Methods), which resulted in approximately 6.2 million mapped reads for *At*-mtRNA (SRA no. SRX4110179) and ∼5.1 million mapped reads for *Bo*-mtRNA (SRA no. SRX4110177) samples. The transcript reads were than compared with the relative genetic loci, *i*.*e*. de novo assembly of the Arabidopsis Col-0 mitogenome (BK010421; (Sloan, *et al*. 2018) (Fig. 1a), and the cauliflower mtDNA established in a collaboration between the lab of Prof. Jeff Mower and our group (Fig. 1b, KJ820683) (Grewe, *et al*. 2014).

**Figure 1.**
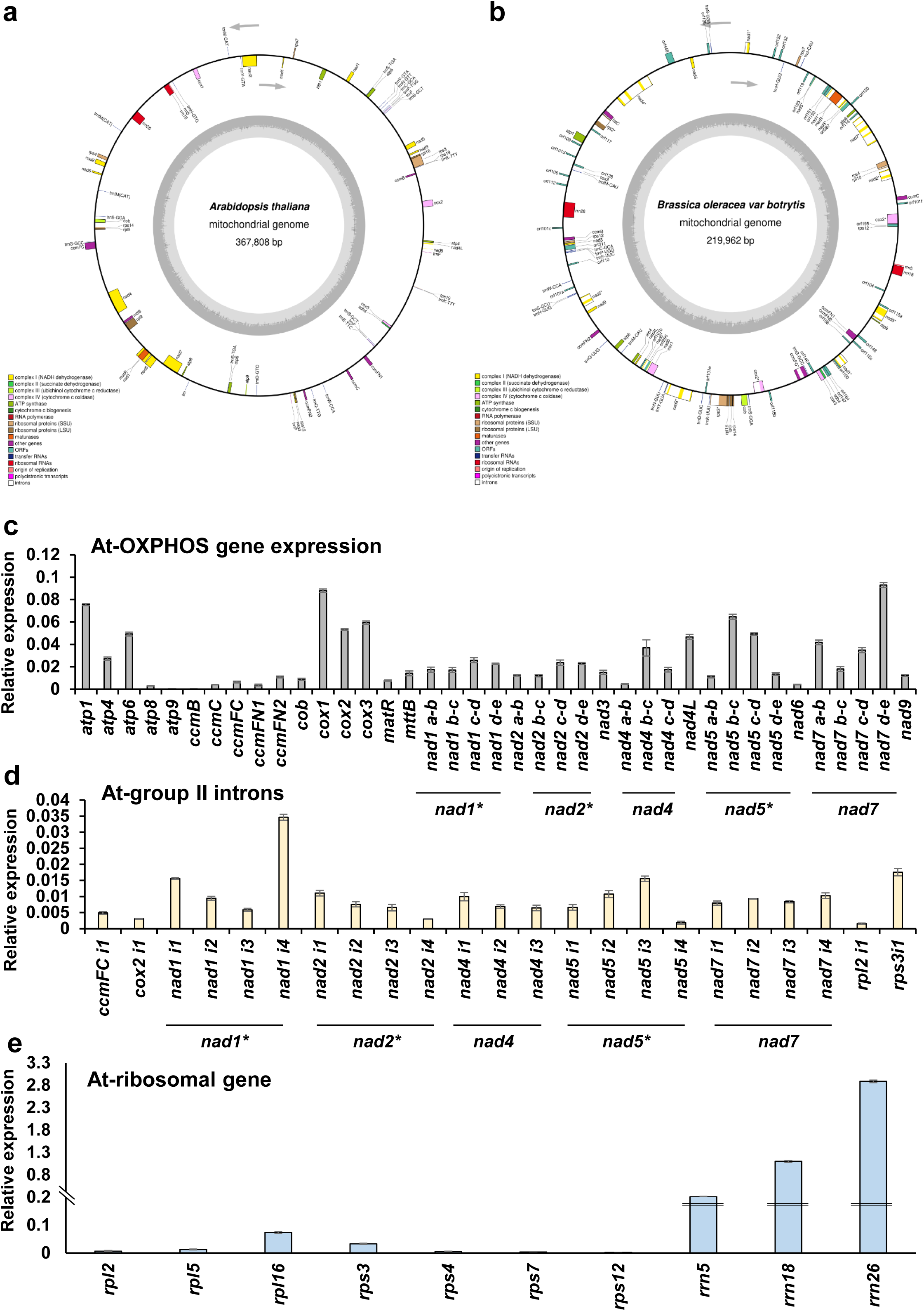
Global expression profiles of *Arabidopsis thaliana* (var. Col-0) and *Brassica oleracea* (var. botrytis) mitochondria. Physical map of (a) Arabidopsis (BK010421) (Sloan, *et al*. 2018) and (b) cauliflower (KJ820683) (Grewe, *et al*. 2014) mitogenomes. Boxes on the inside and outside of the outer circle represent genes transcribed clockwise and anti-clockwise, respectively. The location of each gene is depicted using the systematic name. Gene colors correspond to the functional categories are listed in the legend, while open boxed regions indicate group II intron regions. The figure was created using OGDraw v1.2 (Lohse *et al*. 2013). RNA extracted from 3-week-old seedlings of Arabidopsis plants was reverse-transcribed, and the relative steady-state levels of cDNAs corresponding to the different organellar transcripts were evaluated by qPCR with primers which specifically amplified mRNAs (Sultan, *et al*. 2016a). Histograms showing the relative transcript levels (*i*.*e*. log2 ratios) of (a) OXPHOS genes, (b) pre-mRNAs and (c) ribosomal proteins and rRNAs, after normalization to the *GAPDH* (AT1G13440), *ACTINE2* (At3g1878), *18S-rRNA* (At3g41768), *RRN26* (*i*.*e*. mitochondrial 26S-rRNA, Atmg00020), *RRN5* (Atmg01380) and *RRN18* (Atmg01390) genes. Transcripts analyzed in these assays include the NADH dehydrogenase (CI) subunits *nad1* exons a-b, b-c, c-d, d-e, *nad2* exons a-b, b-c, c-d, d-e, *nad3, nad4* exons a-b, b-c, c-d, *nad4L, nad5* exons a-b, b-c, c-d, d-e, *nad6, nad7* exons a-b, b-c, c-d, d-e, and *nad9*, the complex III *cob* subunit, cytochrome oxidase (complex IV) *cox1, cox2* exons a-b and *cox3* subunits, the ATP synthase (i.e., complex V) subunits *atp1, atp4, atp6, atp8* and *atp9*, genes encoding the cytochrome c maturation (*ccm*) factors, *ccmB, ccmC, ccmFn1, ccmFn2* and *ccmFc* exons a-b, the ribosomal subunits *rpl2* exons a-b, *rps3* exons a-b, *rps4, rps7, rps12, rpl16, rrn26*, and the *mttB* gene. Asterisks indicate to *trans*-spliced transcripts. The values are means of nine reactions corresponding to three biological replicates (error bars indicate one standard deviation).

The Arabidopsis mitogenome (Sloan, *et al*. 2018, Unseld *et al*. 1997) encodes 28 protein-coding genes, 3 rRNAs, 22 tRNAs and a single intronic ORF (the *matR* gene) (Sultan, *et al*. 2016b, Wahleithner *et al*. 1990). In addition, the Arabidopsis mtDNA also harbors almost 500 mtORFs with at least 60 codons and ∼85 mtORFs of >100 amino acids. The mtDNA of cauliflower encodes 54 known genes, *i*.*e*. 33 proteins (including matR), 3 rRNAs and 18 tRNAs) and 35 mtORFs of at least 100 codons (Grewe, *et al*. 2014). Figure 2 indicates the transcriptome landscapes on the physical maps of Arabidopsis and cauliflower mitogenomes. In total, 42 gene clusters (annotated here as Atm’s) representing either *mono*-, *bi*- or *poly-* cistronic transcription units in Arabidopsis (Fig. 2a and Table 1) and 33 transcription units in cauliflower (Fig. 2b and Table 2), were identified by the RNA-seq analyses. The integrity of the RNA-seq data was further supported by RT-PCR analyses (Fig. 3).

**Table 1.**
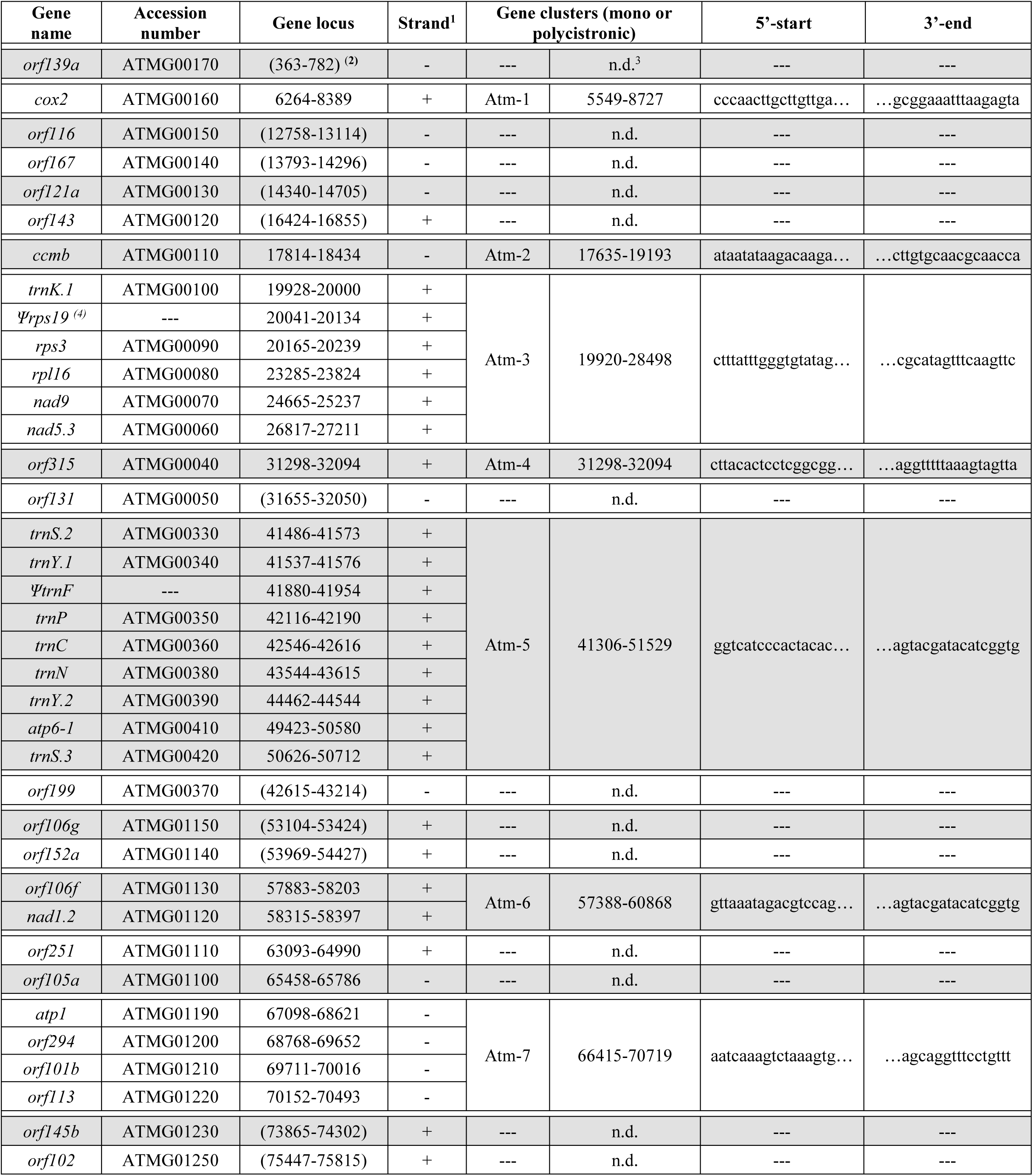
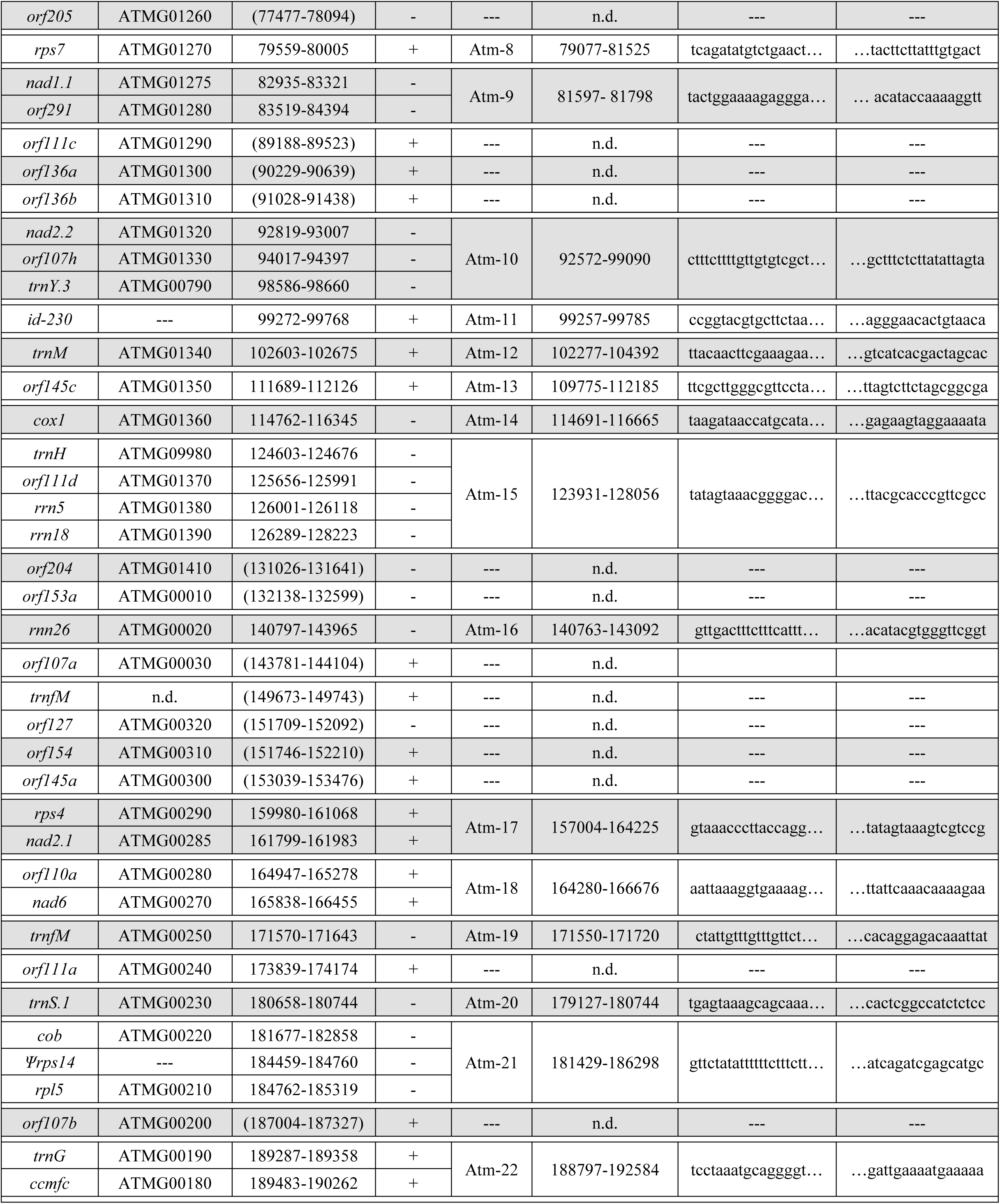
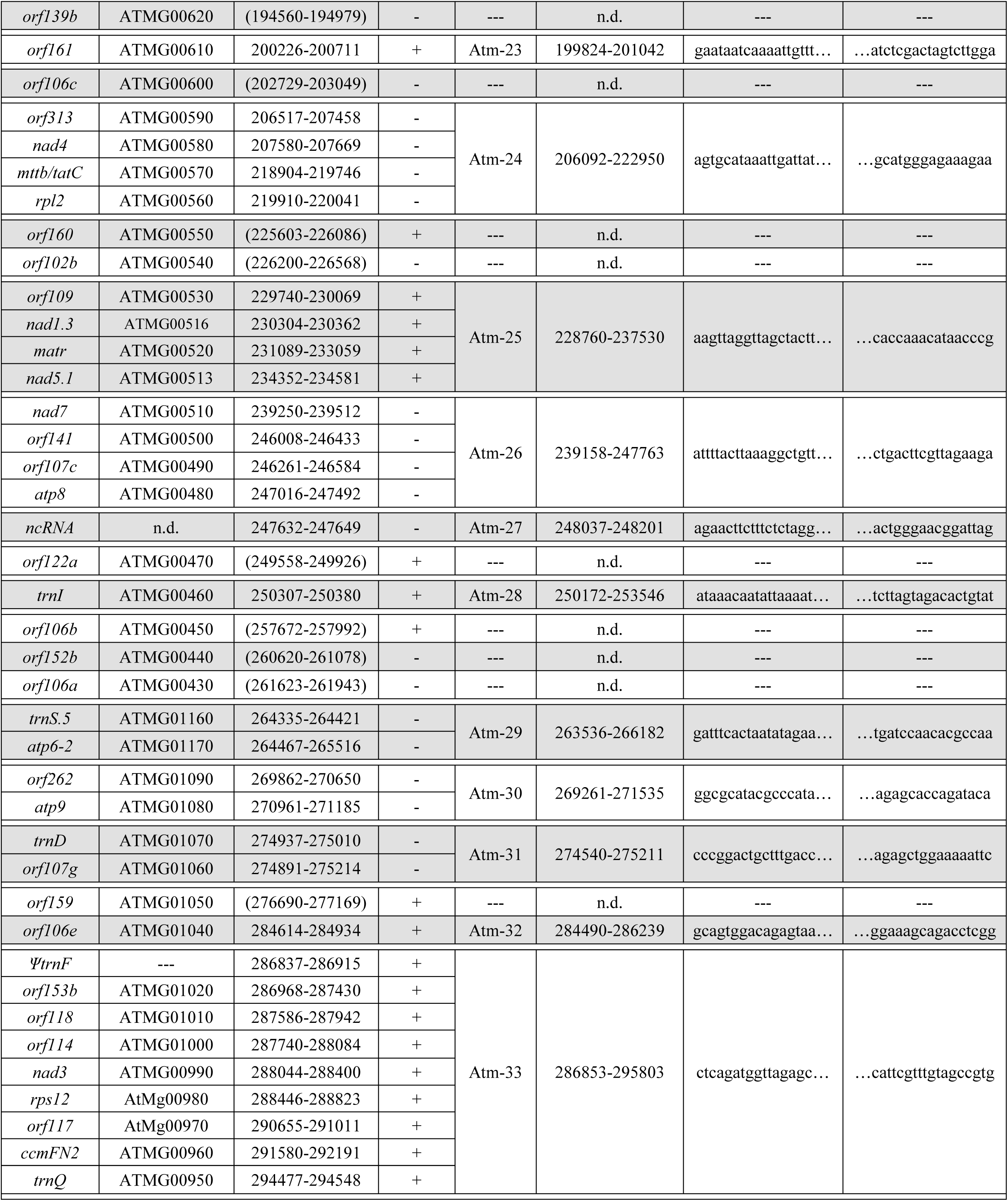
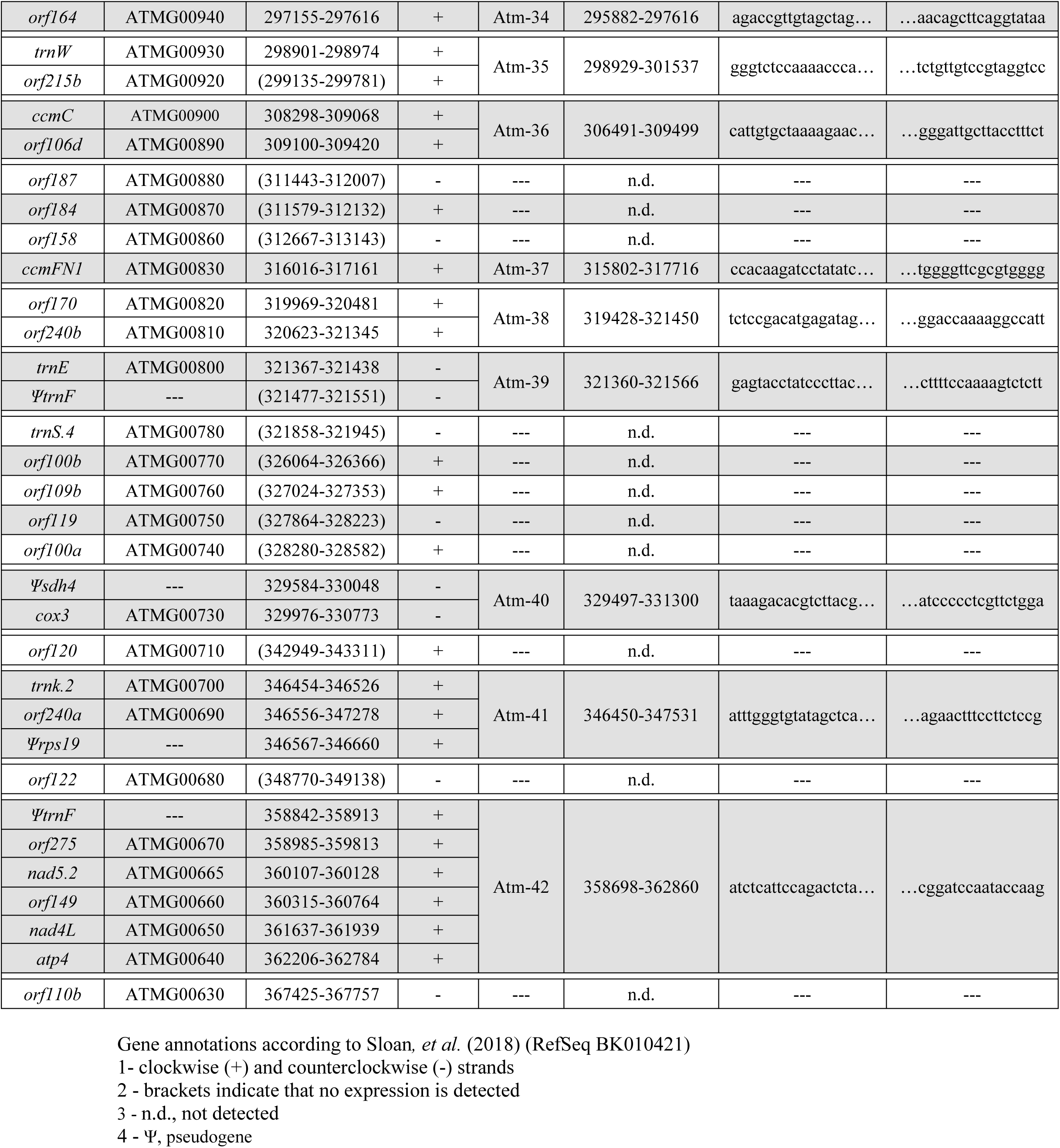
List of Arabidopsis mitochondria gene clusters based on mtRNA-seq analysis.

**Table 2.**
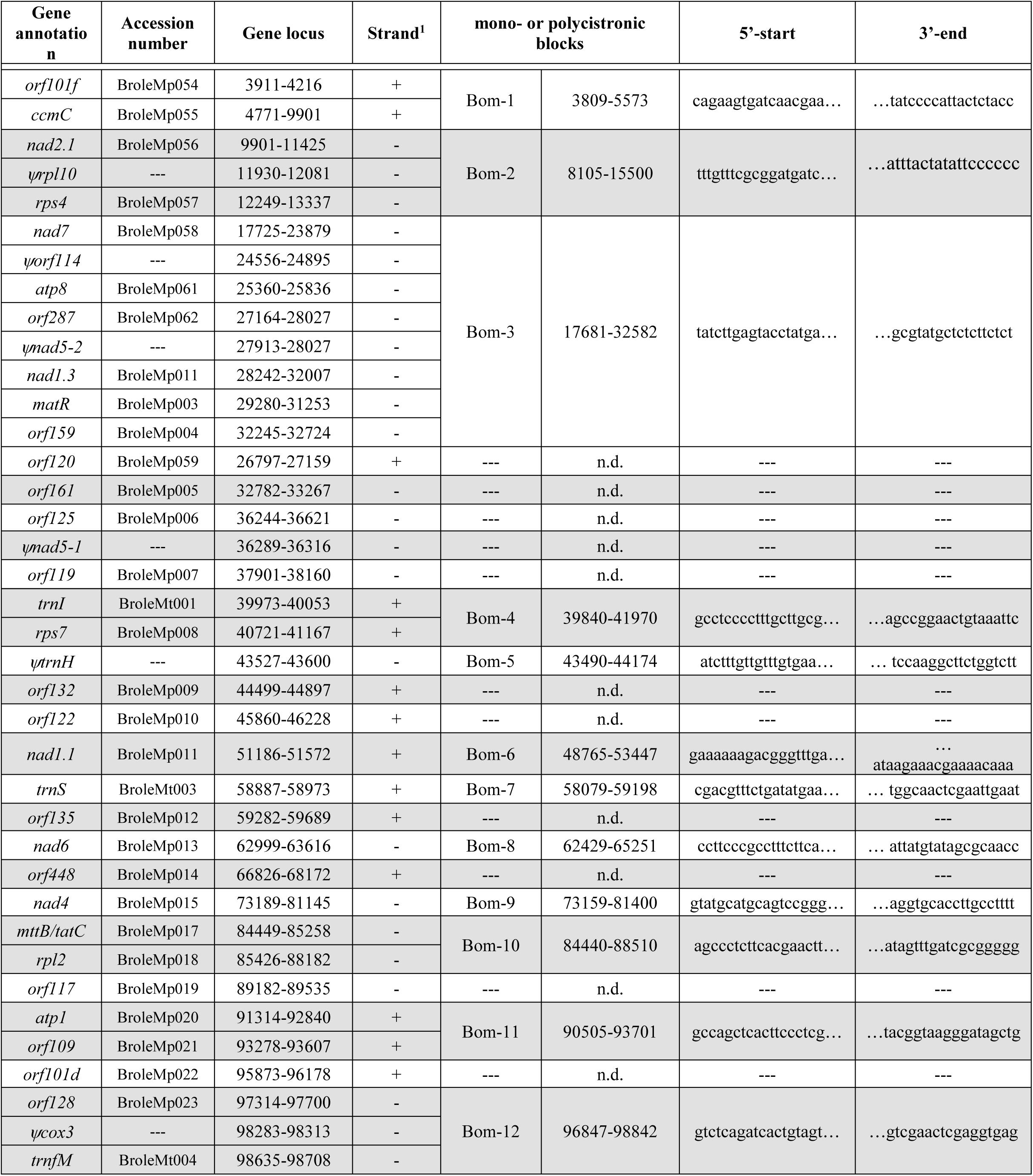
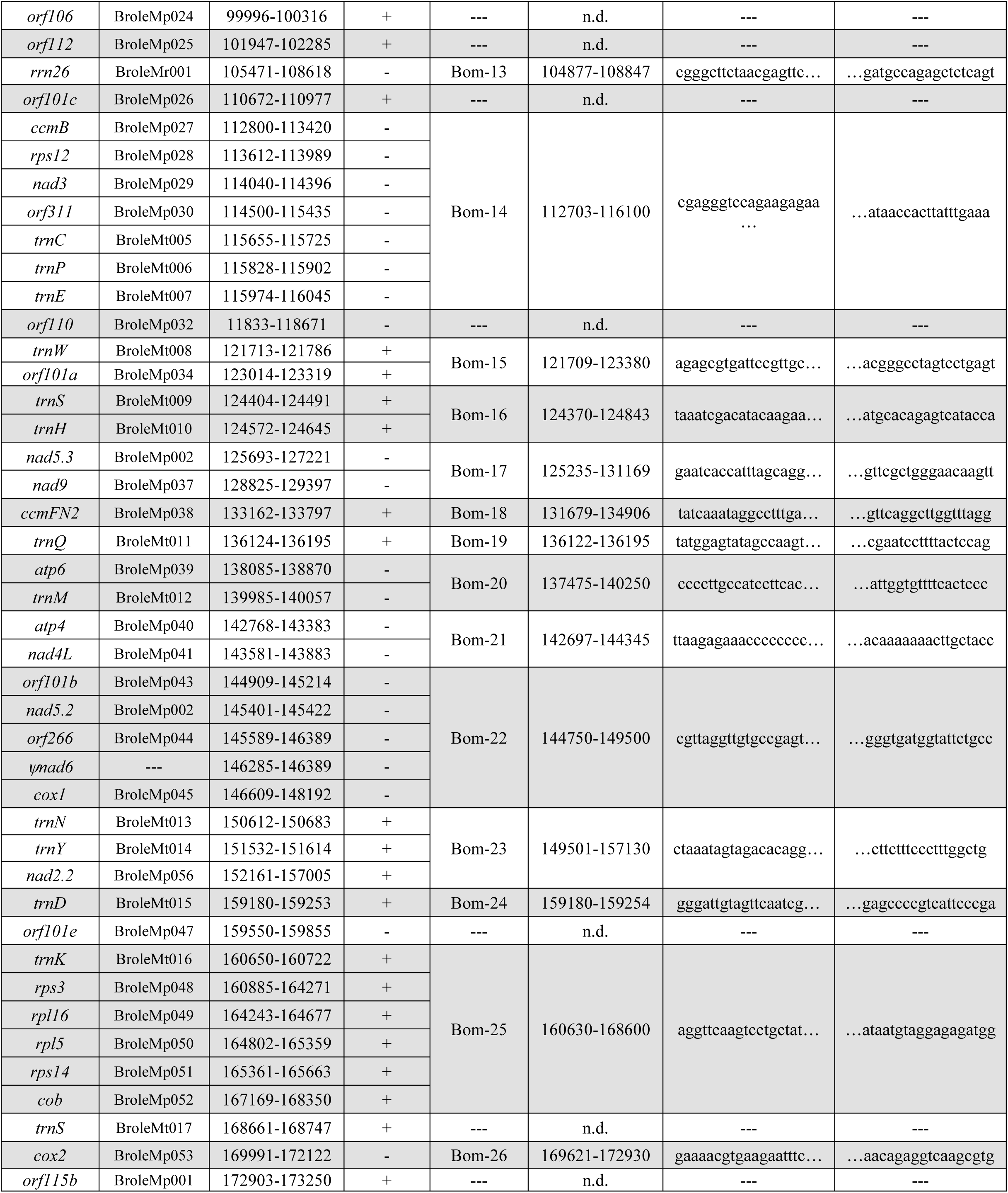
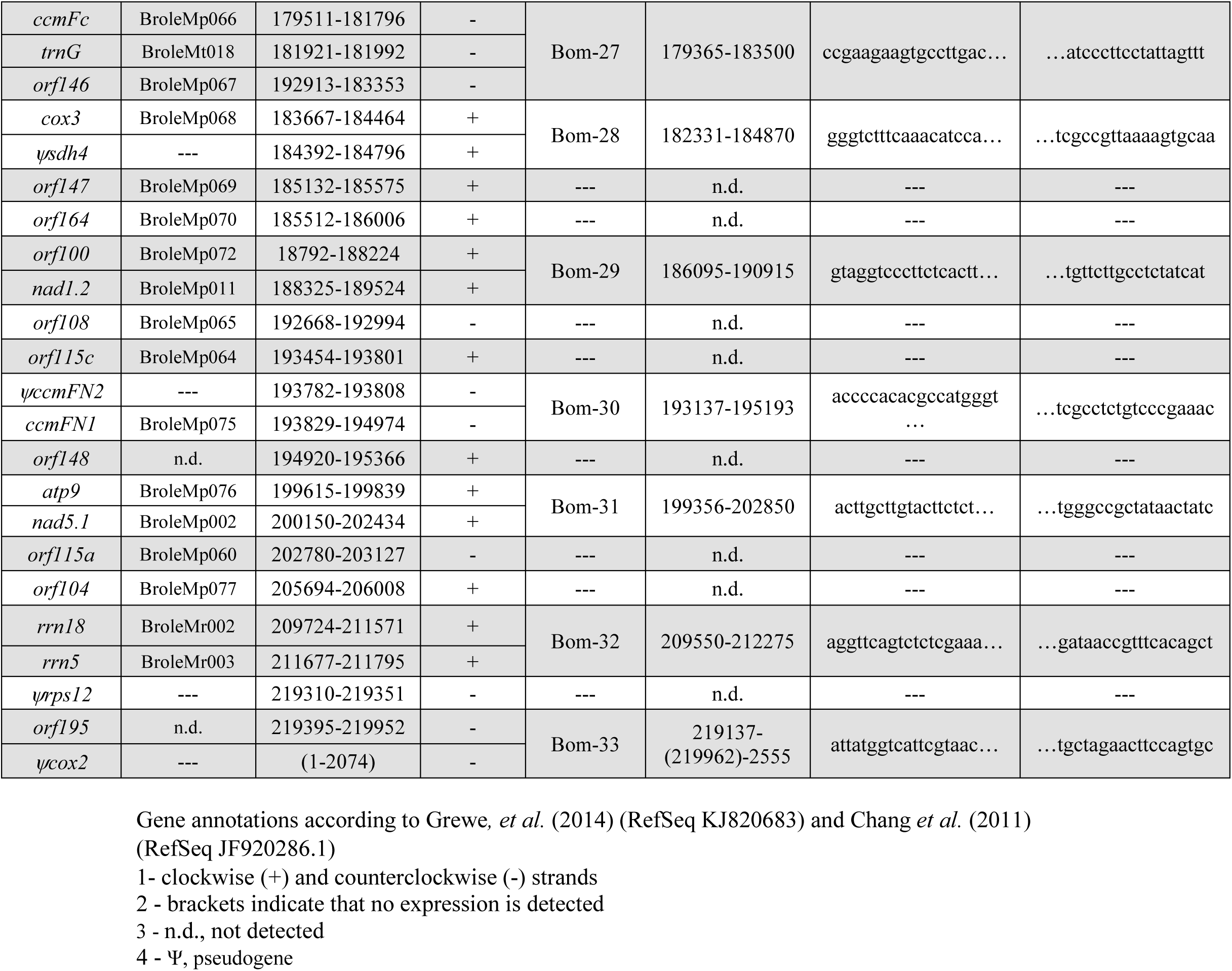
List of cauliflower mitochondria gene clusters based on mtRNA-seq analysis.

**Figure 2.**
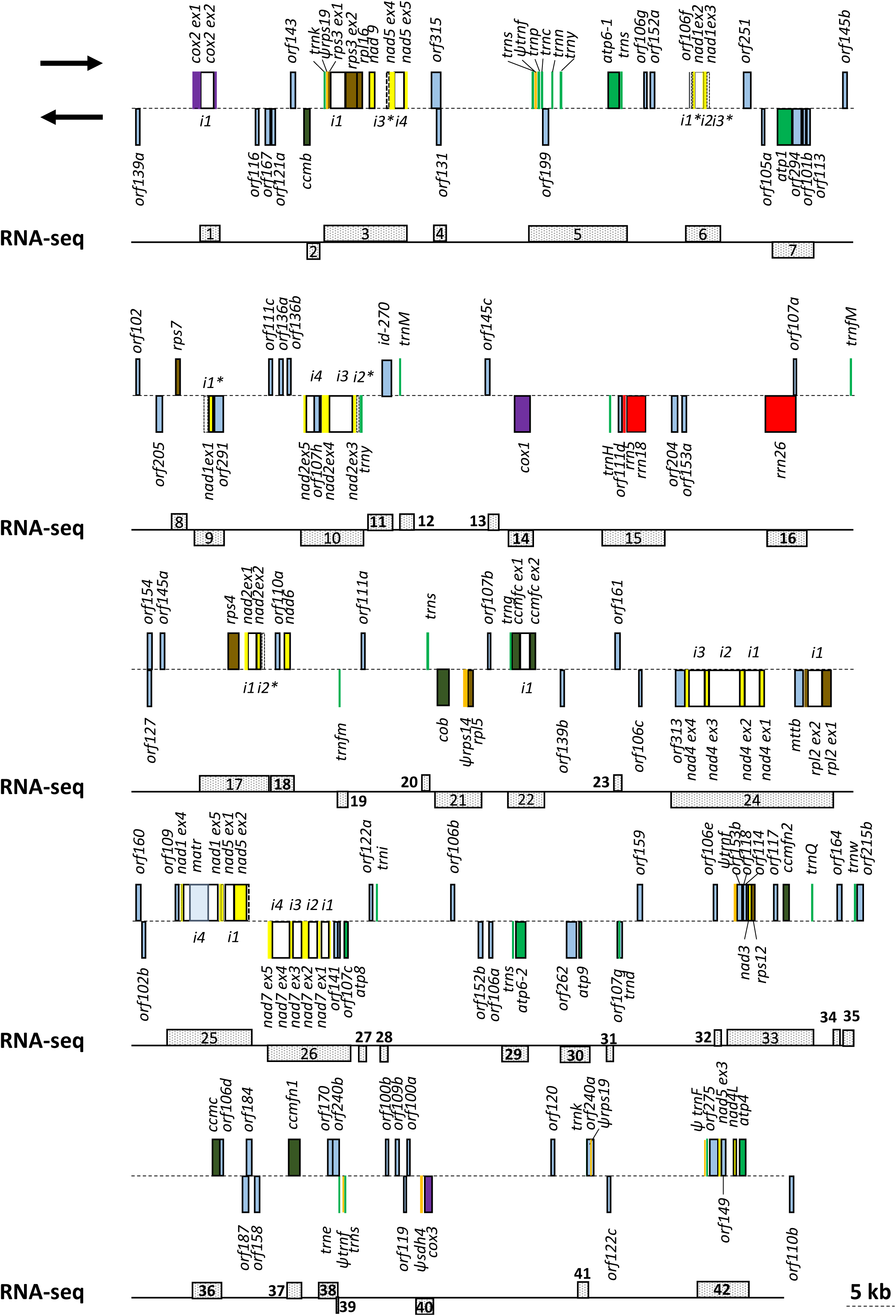
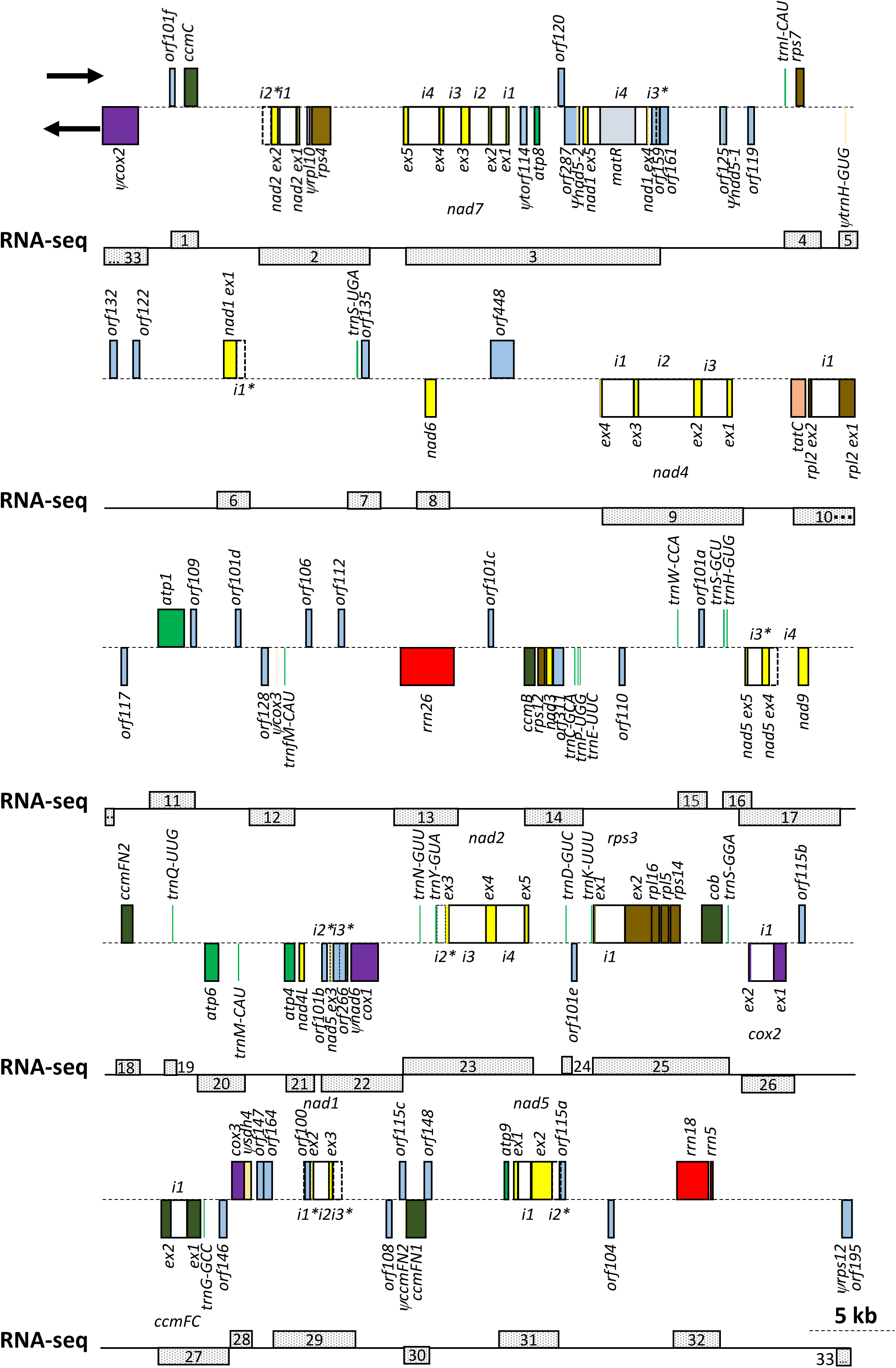
Scheme of the Arabidopsis and cauliflower mitochondria transcriptomes. Overview of the gene clusters of Arabidopsis (a) and cauliflower (b) mitochondria. The gene annotations and loci are indicated in Tables 1 and 2.

**Figure 3.**
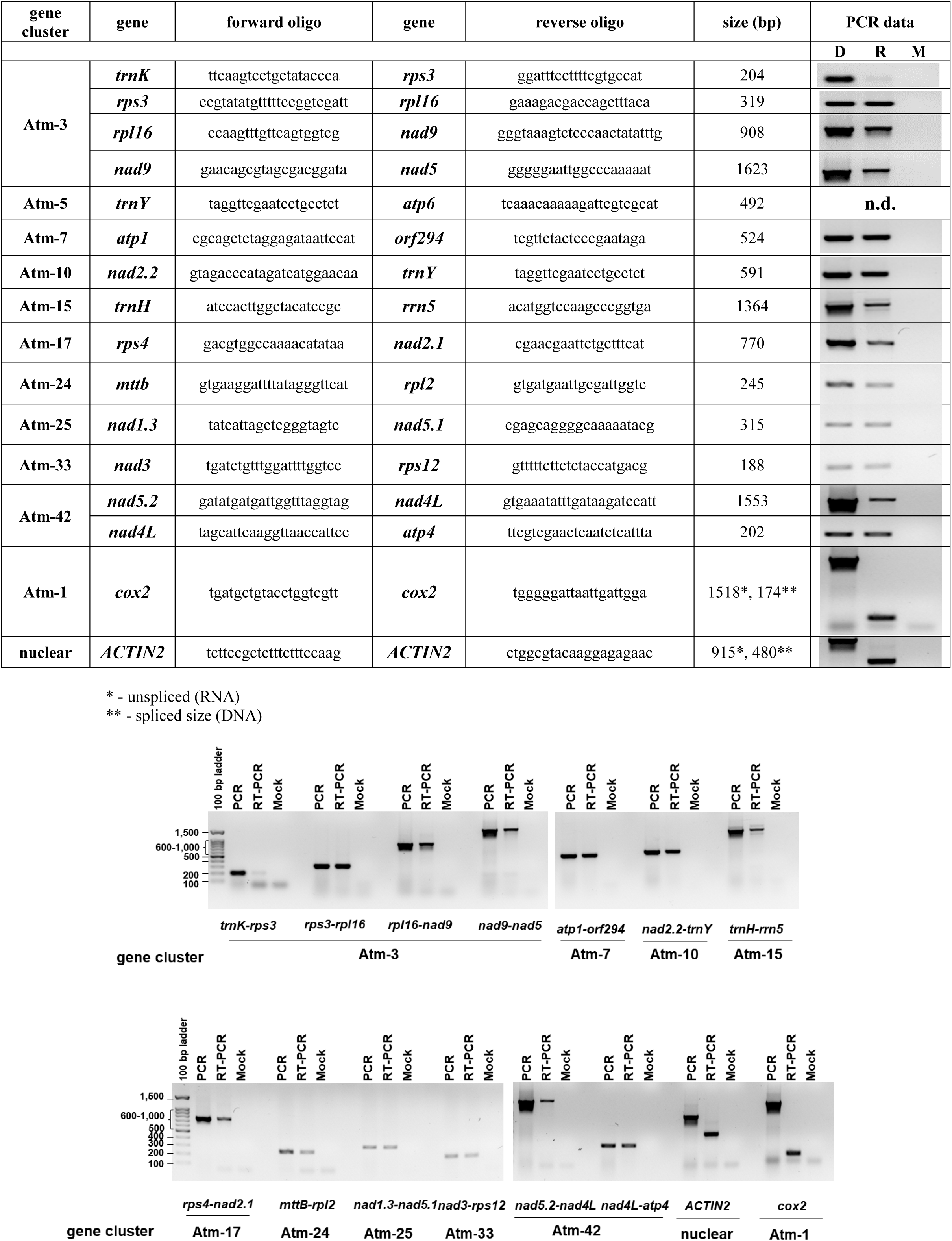
RT-PCR of various mitochondrial gene clusters in *Arabidopsis thaliana* plants. Total DNA and RNA were extracted from 3-week-old seedlings of Arabidopsis plants. The reverse-transcribed At-RNAs and At-DNAs were than used in PCR analyses with oligonucleotides design to junction regions between adjacent genes identified by the RNA-seq as belonging to the same gene-clusters (polycistronic RNAs). To analyze the integrity of the data we used the stand-alone gene of *cox2* and the nuclear gene of ACTIN2 that their genes are interrupted by introns. The splicing yields mature products (the sizes of the unspliced and mature transcripts are indicated).

### Differential expression of mitochondrial genes in *Arabidopsis thaliana* seedlings

Based on microarray, RT-qPCR and RNA-seq data it is suggested that the levels of pre-mRNAs are mainly determined by transcriptional rates, while the accumulation of their corresponding mature transcripts (i.e. mRNAs) is regulated by a balance between transcription and degradation processes (Zeisel *et al*. 2011). Likewise, various studies indicated that the relative accumulation of co-transcribed genes in Arabidopsis mitochondria is only partially, or poorly, related to their expression rates (Binder and Brennicke 2003, Giege *et al*. 2000), further indicating to the complex posttranscriptional events in land plant mitochondria. The relative accumulation (*i*.*e*. steady-state levels) of 3-week-old Arabidopsis seedlings mitochondrial transcripts was analyzed by RT-qPCR. Mapping the Arabidopsis mtRNA-seq reads (SRA n<u>o</u>. SRX4110179) (Murik, *et al*. 2019) to mitochondrial genomic annotation and RT-qPCR data (Figs. 1 and 2) demonstrated differential expression of the 21 mitochondria-encoded genes required for OXPHOS, as well as various ribosomal subunits (Fig 1c-e). These data are perhaps not surprising given that many of the mitochondrial genes in Arabidopsis, unlike their counterparts in the mitochondria of animals, seem to be encoded as *mono*- or bicistronic transcripts (Table 1). Also, we previously indicated the role for the MITOCHONDRIAL TRANSCRIPTION TERMINATION FACTOR 22 (mTERF22) in the regulation of mtDNA expression (Shevtsov *et al*. 2018). mTERF22 affects the expression of numerous mitochondrial genes, besides those of *ccmC, ccmFN1, rpl16, rps3* and *nad5*.*3* genes, which their steady-state RNA levels were not affected by the mutation in *mTERF22* locus (Shevtsov, *et al*. 2018). Interestingly, our RNA-seq data shows that *ccmC* and *ccmFN1* are encoded as mono-cistronic RNAs (Atm-36 and Atm-37, respectively), while the *rpl16, rps3* and *nad5*.*3* genes are all found on the same gene-cluster (Atm-3). The RNA-seq results therefore provide with a molecular explanation to the differences in gene expression in the *mterf22* mutant-lines (Shevtsov, *et al*. 2018).

Complex mechanisms to control gene expression at the transcriptional or translational levels have evolved in all organisms (Hernández *et al*. 2012), allowing them to respond and cope with both developmental and environmental signals. It is anticipated that different subunits belonging to the same protein complex are expressed at equivalent levels. Accordingly, the translation rate of plastidial proteins seems to be closely related with the abundances of their corresponding mRNAs (Chotewutmontri and Barkan 2016). However, the RNA-seq and RT-qPCR data (Figs. 1 and 2) strongly suggest that the levels of various mitochondrial proteins in Angiosperms is regulated at the translational levels. This assumption coincide with differential expression of mtRNAs by ribosomal profiling experiments (Planchard *et al*. 2018), proteomic studies (Huang *et al*. 2020), and the unique properties of the translational apparatus (i.e. mitoribosomes) in Arabidopsis mitochondria (Waltz *et al*. 2019).

The RNA-seq and RT-qPCR also indicated to differential expression of several genes found on the same primary transcript (Fig. 4). These include the clusters of *trnH*-*orf111d*-*rrn5*-*rrn18* (Atm-15), *rps4*-*nad2*.*1* (Atm-17), *cob*-*Ψrps14*-*rpl5* (Atm-24), *orf109*-*nad1*.*3*-*nad5*.*1*(Atm-25), *ΨtrnF*-*orf153b*-*orf118*-*orf114*-*nad3*-*rps12*-*orf117*-*ccmFN2*-*trnQ* (Atm-33) and *ΨtrnF*-*orf275*-*nad5*.*2*-*orf149*-*nad4L*-*atp4* (Atm-42) (Table 1, and Figs. 2-3). For example, we noticed a nearly 7-fold differential expression between *nad3* and *rps12*, about 5-fold between *rps12* and *ccmFN2* (the Atm-33 cluster), and about 4-fold between *rrn5* and *rrn18* genes (the Atm-15 cluster) (Fig. 4b). Differently from the reported influence of attenuated transcription between genes found on the same cistron (Mercer, *et al*. 2011, Turk *et al*. 2013), the levels of mtRNAs is not related to their order on the polycistronic primary transcript. We therefore assume that the differential gene expression patterns in Arabidopsis mitochondria are related to post-transcriptional RNA processing or decay. Taken together, these results correlate with mitochondrial transcriptome data of other eukaryotic cells, showing that co-transcribed mtRNAs have differential abundance. Accordingly, genetic and biochemical data have implicated RBPs in every step of plant mitogenome expression, including transcription, RNA processing and translation (reviewed by *e*.*g*. Zmudjak and Ostersetzer-Biran (2017)).

**Figure 4.**
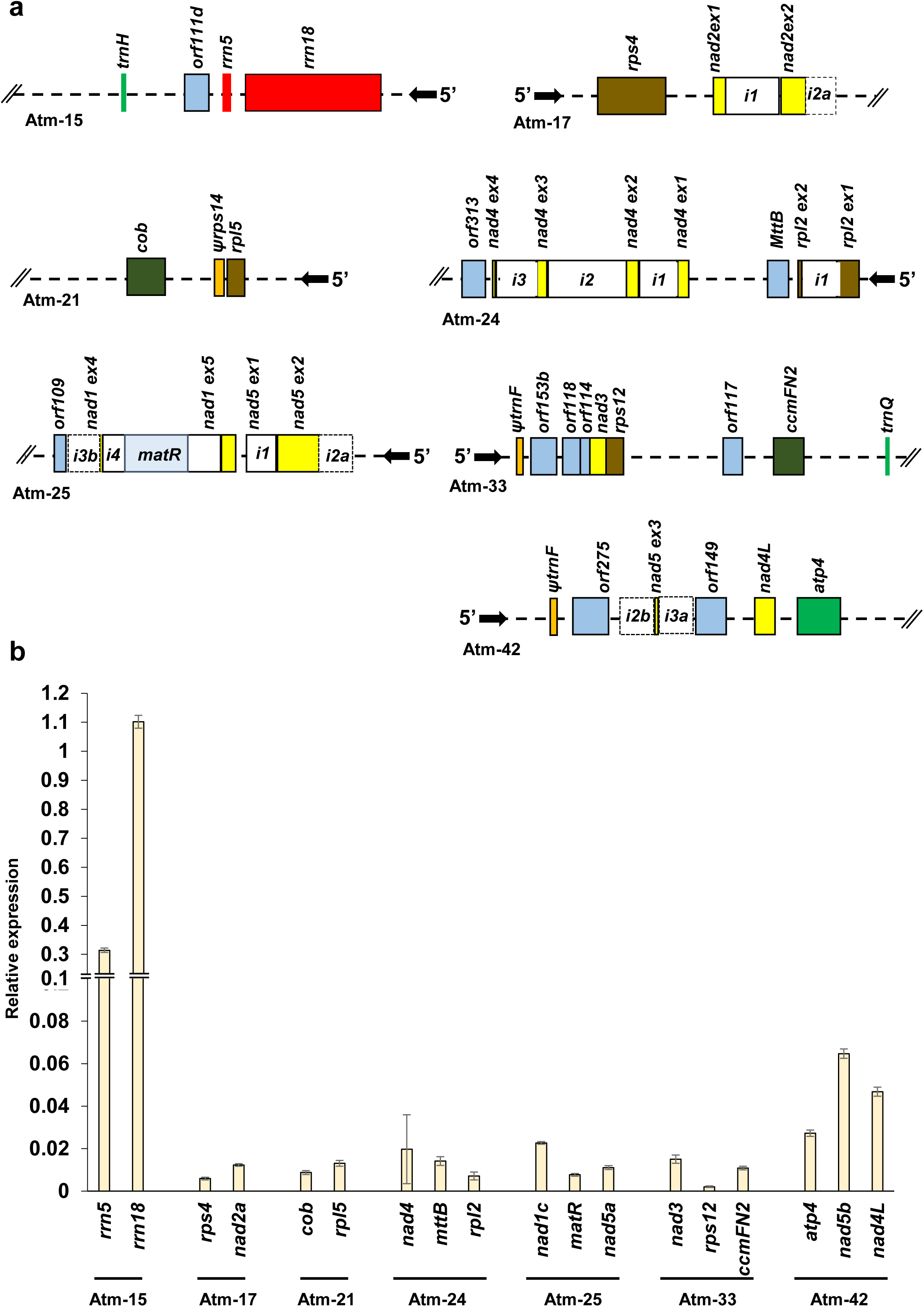
Relative abundances of Arabidopsis genes transcribed by polycistronic pre-RNA transcripts. (a) Overview of several gene clusters in Arabidopsis mitochondria. The gene annotations and loci are indicated in Tables 1 and 2. (b) RNA extracted from 3-week-old seedlings of Arabidopsis plants was reverse-transcribed, and the relative steady-state levels (log2 ratios) of cDNAs corresponding to *rrn5, rrn18, rps4, nad2*.*1, cob, rpl5, nad4, mttB, rpl2, nad1*.*3, matR, nad5*.*1, nad3, rps12, ccmFN2, atp4, nad5*.*2*, and *nad4L* (clusters Atms’ 15, 17, 21, 24, 25, 33 and 42, Table 1) were quantified by RT-qPCR (Sultan, *et al*. 2016a), after normalization to *GAPDH* (AT1G13440), *ACTINE2* (At3g1878), *18S-rRNA* (At3g41768), *rrn26* (*i*.*e*. mitochondrial 26S-rRNA, Atmg00020), *rrn5* (Atmg01380) and *rrn18* (Atmg01390) genes. The expression of *rrn18* is arbitrarily set to 1.0. The values are means of nine reactions corresponding to three biological replicates (error bars indicate one standard deviation).

### Analyses of the 5’ and 3’ ends of mitochondrial RNAs of Arabidopsis and cauliflower

The trimming of precursor transcripts is an essential step in mtRNA maturation (see e.g. (Binder *et al*. 2013, Forner *et al*. 2007, Zmudjak and Ostersetzer-Biran 2017). These activities are generally accomplished by the activities of various endoribonucleases and exoribonucleases. In mitochondria of animals, RNA stem-loop or tRNA-like structures seem to be important for stabilizing the transcripts from excess degradation. Analysis of the mRNA extremities of the mitochondrial encoded protein-coding genes in *Arabidopsis thaliana* plants revealed to RNA stem-loop structures (termed as t-elements) at the 5’ or 3’ termini of some mRNAs (Dombrowski *et al*. 1997, Forner, *et al*. 2007). However, the vast majority of the organellar transcripts in plants seem to lack similar structural elements at their postulated 5’- or 3’-ends (Forner, *et al*. 2007). Alternatively, RBPs may have acquired similar functions in the organelles of land plants to those of stem-loop regions, most likely by blocking 5′-to-3′ or 3′-to-5′ RNA exonucleolytic degradation, and thereby defining the transcript ends (Small *et al*. 2013). Pentatricopeptide repeat (PPR) proteins play pivotal roles in these activities (Barkan and Small 2014). RNA-seq data of small organellar RNAs supported this hypothesis for the maturation of plastidial transcripts in Maize (Hotto *et al*. 2011, Ruwe and Schmitz-Linneweber 2012, Zhelyazkova *et al*. 2012), but a mitochondria-wide gene expression related for the process in angiosperms is still awaiting investigation. Small RNAs that overlap with the processed 3′-ends, but not with the 5’-ends, of various mtRNAs was noted in Arabidopsis (Ruwe *et al*. 2016).

Sequencing the mitochondrial transcriptome provided us with more insights into the processing of 5’ and 3’ termini of angiosperms mitochondrial transcripts. This was made possible by relative read counts of *Arabidopsis thaliana* and *Brassica oleracea* mtRNA-seq data. Tables 1 and 2 show the gene clusters and 5’ and 3’ termini of the mitochondrial genes, including tRNAs, rRNAs, coding-regions, as well as exon and intron sequences, and all the annotated mtORFs in Arabidopsis and cauliflower mitochondria, respectively. We further compared the RNA-seq data with the proposed 5’ and 3’ termini of various Arabidopsis transcripts analyzed by circular RT-PCR (cRT) assays (Forner, *et al*. 2007). In general, our RNA-seq data, corresponding to genes expressed as mono- or bicistronic RNAs, was in a good agreement with the cRT experiments (Fig 5). We assume that differences in the processing sites seen between the RNA-seq and cRT data, are attributed to the different methodologies used in the assays, *i*.*e*. RT followed by cDNA sequencing in Forner, *et al*. (2007) *vs*. the global mtRNA-seq data, or to differences in gene annotation (here we use the de novo assembly of the Arabidopsis Col-0 mitogenome, *i*.*e*. accession no. BK01042) (Sloan, *et al*. 2018). More notable differences in mtRNA termini sites were observed in genes transcribed in large polycistronic RNAs and particularly in cases of ‘split-genes’, separated by *trans*-spliced introns (i.e. *nad1, nad2, nad5* genes) (Figs. 1-3, 5, and Table 1).

**Figure 5.**
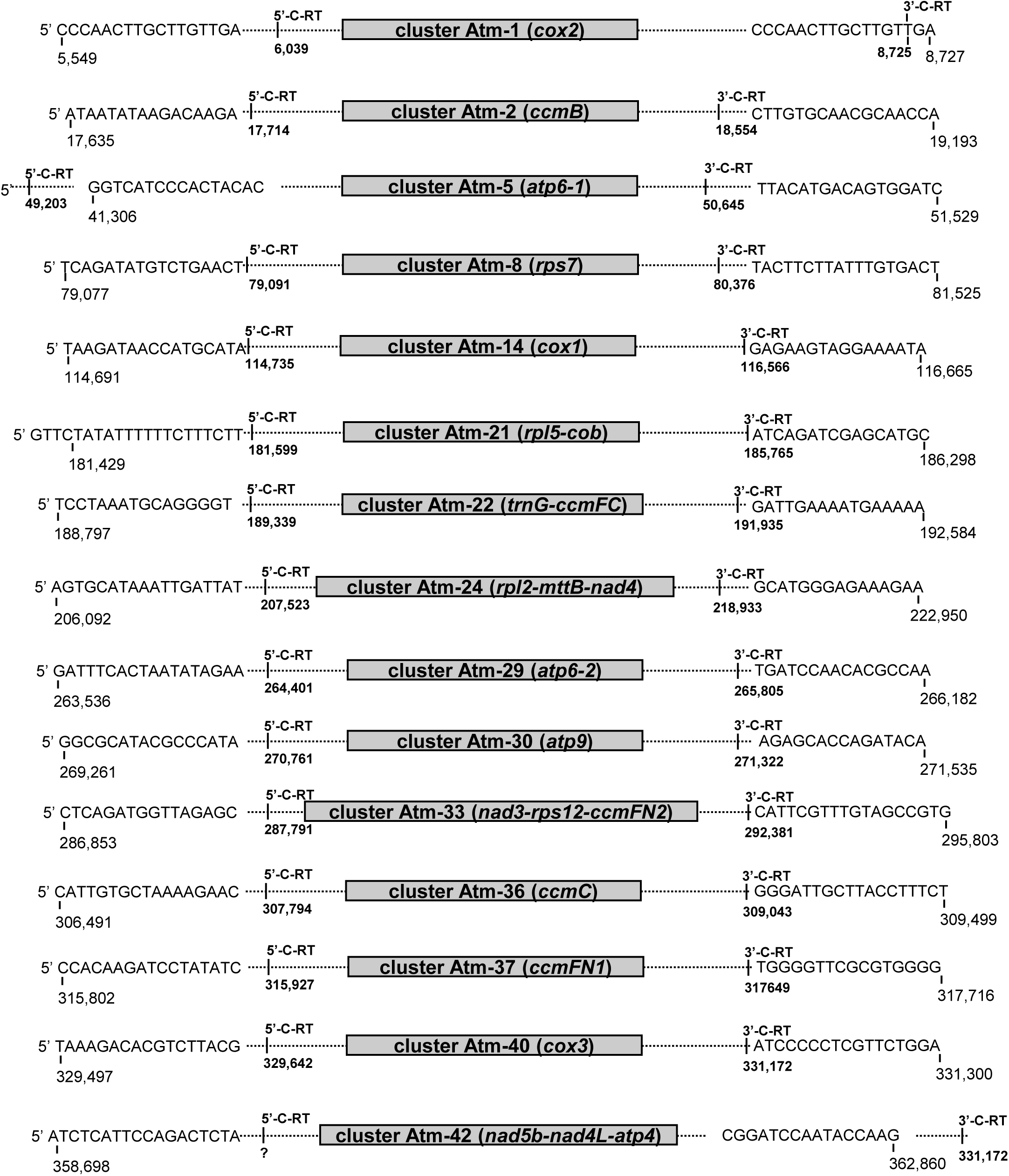
Arabidopsis 5’ and 3’ termini as determined by mtRNA-seq data. The scheme in the figure depicts the organization of various genes in the mitochondrial DNA of *Arabidopsis thaliana*. Details of the 15 different sequences obtained from the RNA-seq data are shown. The positions of the 5’ and 3’ termini of the set of genes as obtained by cRT experiments (Forner, *et al*. 2007) are indicated in the scheme.

In Arabidopsis and cauliflower mitochondria, the maturation of *nad1* involves the joining of five exons, encoded by three individually-expressed gene fragments (*i*.*e. nad1*.*1, nad1*.*2* and *nad1*.*3*, Atm’s 9, 6 and 25, respectively), which are spliced through two *trans*-(introns 1 and 3) and two *cis*-(introns 2 and 4) events. Likewise, the maturation of *nad2* involves the splicing of 3 *cis*-(i.e. *nad2* introns 1, 3 and4) and one *trans*-spliced intron (*nad2* i2) found on two individual transcripts (*nad2*.*1* and *nad2*.*2*, Atm’s 17 and 10, respectively), whereas the maturation of *nad5* involves the joining of five exons, encoded by three individually-expressed gene fragments (*i*.*e. nad5*.*1, nad5*.*2* and *nad5*.*3*, Atm’s 25, 42 and 3, respectively), where *nad5* introns 2 and 3 are physically separated on the mitogenome and are spliced in trans (Figs. 1-3, 5, and Table 1). It is expected that the cRT methodology is not an appropriate tool to address the 5’ or 3’ ends of polycistronic RNAs of split-genes, which their exons are physically separated on the mitogenome.

### The postulated roles of RBPs in 5’ and 3’ processing of angiosperms mtRNAs

No significant homology or conserved sequence motifs could be identified in the postulated 5’or 3’ ends of the mtRNAs in Arabidopsis or cauliflower (Fig 6). We hypothesize that the lack of consensus regions is due to the roles of different RBPs, as PPR factors (Barkan and Small 2014), which bind to specific regions and play central roles in determining the 5’ and 3’ termini of mitochondrial transcripts in plants. A question remains is to whether such an assumption can be supported by *in vivo* studies. To address this, we compared the mtRNA-seq data with Arabidopsis mutants affected in the processing of mitochondrial transcripts.

**Figure 6.**
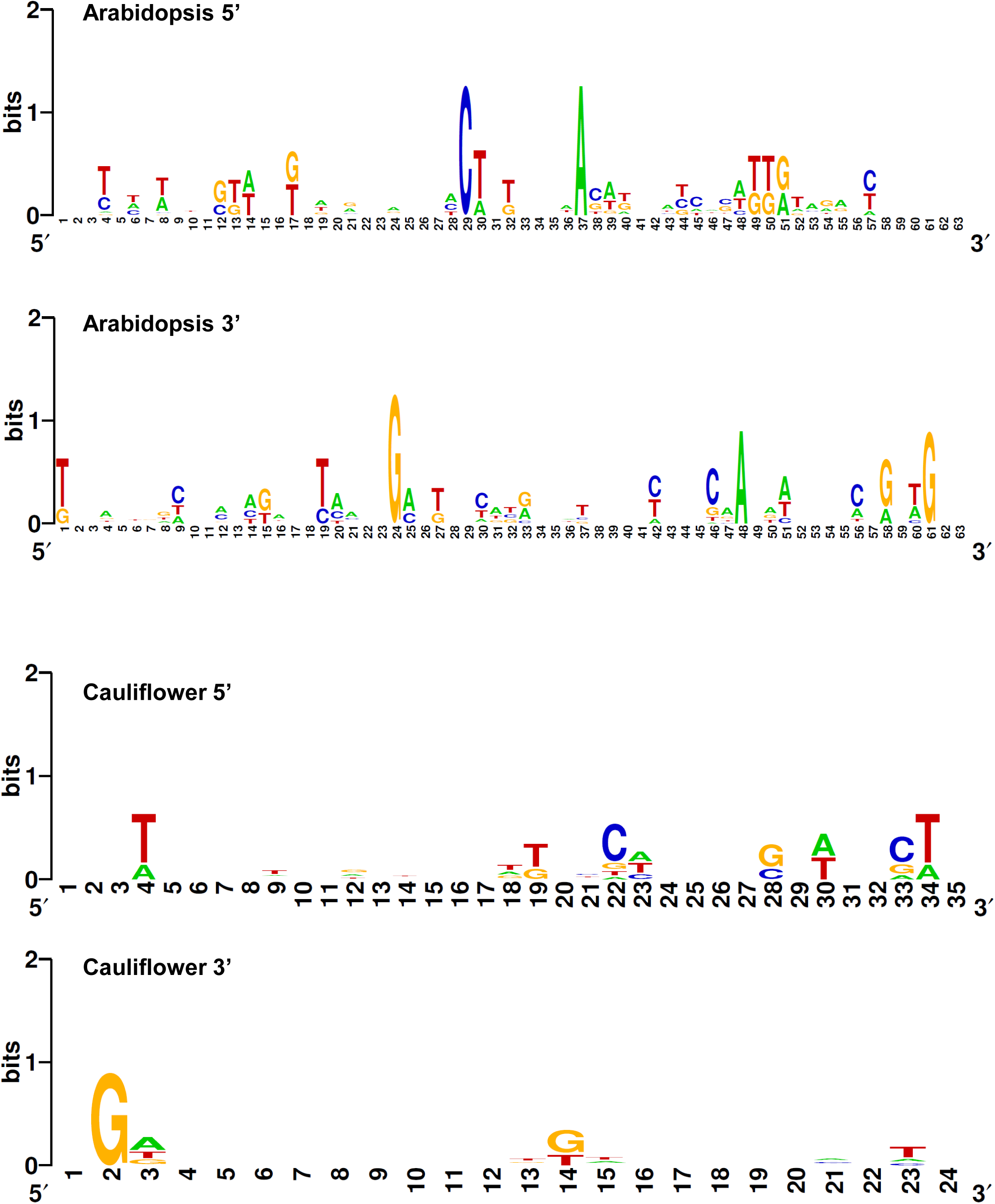
Common termini patterns in Arabidopsis and cauliflower mitochondria. The identity of putative consensus sequence motifs in the 5’ or 3’ termini of Arabidopsis and cauliflower mtRNA, was analyzed by the HOMER (Heinz *et al*. 2010) and MEME (Bailey *et al*. 2009) programs.

The so-called ‘footprints’ of only a few mitochondrial factors has been experimentally verified to date. The PPR-related MTSF1 protein was shown to be associated with the 3’ UTR of *NADH dehydrogenase 4* (*nad4*) and thereby stabilizes its RNA transcript (Haili *et al*. 2013). In fact, the postulated binding site of MTSF1 (5’-CAAAGAAAAGAAAAACGGGU) is mapped at the 3’ terminus of Atm-24 cluster, encoding the *rpl2, mttB* and *nad4* genes (Table 1, and Fig. 5). The PPR MTSF2 protein was shown to stabilizes the 3’-end of *nad1*.*1* precursor transcript (Wang *et al*. 2017). The postulated binding site of MTSF2 (*i*.*e*. 5’-<u>GAUACAUCGGUG</u>UAAAAGGU) (Wang, *et al*. 2017) is located within the predicted 3’ termini (5’-AGUACGAUACAUCGGUG, nts 57,388-60,868) of the Atm-6 cluster, encoding the *nad1*.*2* transcript (Table 1). Likewise, the predicted binding site (*i*.*e*. 5’-<u>AUACAUCGGUG</u>UAAA) of another PPR factor that PPR19, which is also required for the stabilization of *nad1*.*2* transcript (Lee *et al*. 2017) is found at the 3’-end of *nad1*.*2*. RPF7 is another PPR factors that was shown to act in the maturation of the 5′ terminus of *nad2* mRNA (Stoll *et al*. 2014). However, the binding site of PRF7 has not been deciphered yet. We further used the RNA-seq data to analyze the putative binding site of the PPR MSP1025/EMB1025 protein (Best *et al*. 2019). Notably, the predicted RNA binding site of MSP1025 is mapped at the very 3’-end of *nad1*.*1* (*i*.*e*. 5’-ACAUACCAAAAGGUU, nucleotides 81597-81798) (Table 1-Atm-9 gene cluster, and Best et al. MS submitted). These data further support the integrity of the mtRNA-seq data.

### The expression of tRNA genes in Arabidopsis mitochondria

Most mitochondria contain their own genetic system (mtDNA, mitogenome), with an intrinsic protein-synthesis machinery. Transfer RNAs (tRNAs) serve as adaptor molecules, linking a codon in an mRNA with the corresponding amino acid. The mitochondrial translation system thus requires a full set of tRNAs, to enable the incorporation of the 20 proteinogenic amino acids into the nascent polypeptide chain. The pool of tRNAs in vascular plant mitochondria consists of both native (*i*.*e*. mitochondria-encoded) and imported tRNAs (Maréchal-Drouard *et al*. 1999, Salinas-Giegé *et al*. 2015). The mitochondrial genomes of Arabidopsis and cauliflower encode all but six amino acids (Grewe, *et al*. 2014, Sloan, *et al*. 2018, Unseld, *et al*. 1997). Thus, the import of cytosolic tRNA into the organelles is a prerequisite for plant mitochondrial translational (Salinas *et al*. 2008). The mitogenome of Arabidopsis encodes 28 tRNAs (Sloan, *et al*. 2018, Unseld, *et al*. 1997). These include two *trnK* and two *trnfM* genes, three *trnY*, five *trnS*, the *trnC, trnD, trnE, trnG, trnH, trnI, trnM, trnN, trnP, trnQ, trnW* genes and four pseudo-*trnF* (*ΨtrnF*) loci. Our RNA-seq data suggest that 8 of these are clustered together (*i*.*e*. the Atm-5 gene-cluster), and confirmed the expression of many of the tRNA genes (Fig. 2 and Table 1). Out of the two *trnfM* genes, an expression was supported for only a single gene (cluster Atm-19) (Table 1). The expression of *trnW* was close to the limit of detection, while no expression was confirmed for the four *ΨtrnF* genes (Table 1). These data seem to coincide with previous reports indicating the import of a cytosolic *trnF* (Chen *et al*. 1997), and the lack of expression of *trnW* in the mitochondria of Arabidopsis plants as evident by RT-PCR (Duchêne and Maréchal-Drouard 2001).

### The processing of group II introns in Brassicales mitochondria

RNA splicing is an essential post-transcriptional processing step in the maturation of numerous nuclear and organellar introns in plants, where intervening non-coding sequences (introns) are removed, and the coding sequences (exons) are ligated together to form a functional mRNA transcript. Group II introns are large catalytic RNAs, found in the genomes of many prokaryotes and the organellar of some lower eukaryotes, but are particularly prevalent in the organelles of terrestrial plants (Bonen 2008). In angiosperm’s mitochondria, group II introns reside in the sequences of many critical protein coding genes and their excision is therefore essential for organellar biogenesis and respiratory-mediated functions (Brown, *et al*. 2014, Schmitz-Linneweber, *et al*. 2015, Zmudjak and Ostersetzer-Biran 2017). Their splicing is facilitated by numerous protein cofactors, which belong do diverse sets of RBPs, most of which are encoded in the nucleus, *e*.*g*., maturases, RNA-helicases, mTERFs, PORR and PPR related proteins (Brown, *et al*. 2014, Colas des Francs-Small and Small 2014, Schmitz-Linneweber, *et al*. 2015). Figure 7 shows the RNA-seq data corresponding to the 23 introns in Arabidopsis mitochondria (Sloan, *et al*. 2018, Unseld, *et al*. 1997).

**Figure 7.**
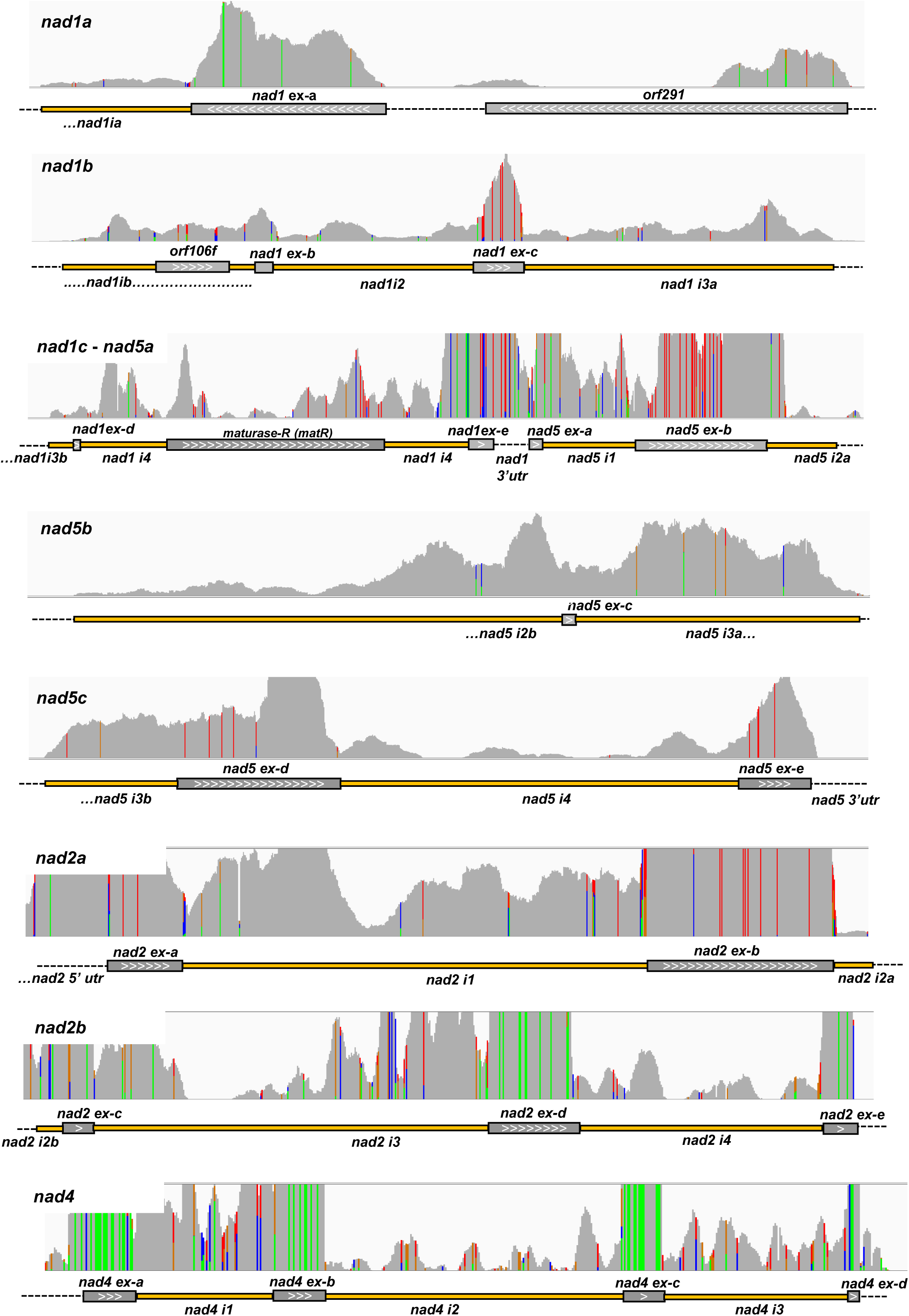
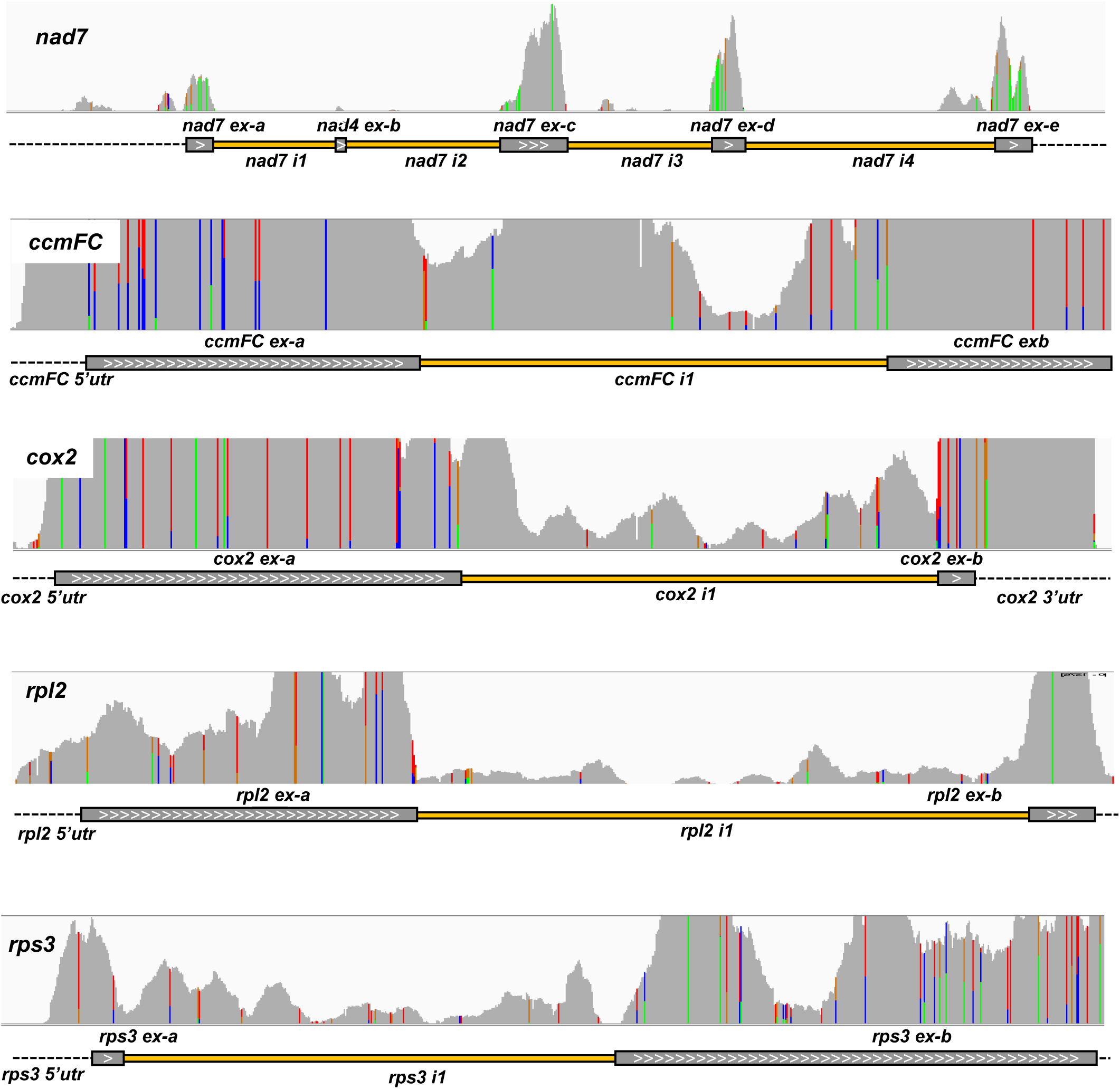
Representative tracks of the 23 group II introns in Arabidopsis mitochondria. Integrative Genomics Viewer (IGV) browser was used to indicate the mtRNA-seq data peaks, corresponding to the 23 group II introns in Arabidopsis mitochondria. For each panel, grey track is the input signal, whereas green, blue or red indicate to mtRNA base changes vs the input mtDNA Sloan, *et al*. (2018) (RefSeq BK010421). The RefSeq gene map is presented in at the bottom of each panel showing the overall gene structure.

The RNAs-seq and RT-qPCR data indicate that many group II introns, as well as their pre-RNAs, in Arabidopsis mitochondria are accumulating to high levels (Figs. 1 and 7). In some cases (*e*.*g. nad1* introns 2 and 3, *ccmFc* i1, *nad4* i1, and *nad5* i2 and i3) their levels seem equivalent to those of their corresponding exonic regions (Figs. 1 and 7). Notably, the steady-state levels of partially-matured RNAs belonging to *trans*-spliced *nad1, nad2* and *nad5* genes were stoichiometry uneven, as apparent by RT-qPCR analyses (Fig. 1) and mtRNA-seq data (Fig. 7). It remains unclear whether the partially processed transcripts found in lower abundances are a limiting factor in the maturation of their corresponding organellar mRNAs. Nonetheless, the accumulation of released intron sequences and their pre-RNAs in the mitochondria of Arabidopsis coincide with previous reports showing that organellar group II introns in plants are generally more stable than their spliceosomal counterparts (Barkan 1989, Li-Pook-Than and Bonen 2006). Accordingly, accumulation of unspliced pre-RNAs is noted during early plant development in both the plastids and mitochondria (Barkan 1989, Li-Pook-Than *et al*. 2004), strongly suggesting a tissue or developmental regulated intron splicing. Ribosomal profiling further indicated that intron-containing precursor transcripts accumulate to high levels in *Zea mays* (corn) plastids, some of which were shown to be associated with stromal ribosomes (Zoschke and Barkan 2015, Zoschke *et al*. 2013). Yet, it remains unclear whether these pre-RNAs are functionally translated, or whether the splicing of plastidial introns might be concurred with the translation reaction (or assisted by specific ribosomal cofactors). Ribosomal footprinting of Arabidopsis mitochondria (Planchard, *et al*. 2018) indicated that UTR and intron sequences are only poorly represented in the organellar polysome fractions.

Following the splicing reaction, the spliceosomal introns are degraded rapidly (with a half-life of only a few seconds), and typically exist in very low or below detectable levels, in vivo (Sharp 1987). The biological imperatives for the turn-over of the released introns include: (a) the recycle of ribonucleotides, and (b) to free their associated RBPs for sequential rounds of transcription and splicing. Although the majority of spliceosomal intron lariats are degraded following pre-RNA splicing, some of the introns are more stable and persist post-splicing (Taggart and Fairbrother 2018). These are hypothesized to have additional post-splicing functions. In would be interesting to study whether the accumulation of free group II intron lariats has a biological significance. In this respect, comparing the mtDNA sequences of different plants indicated the loss of group II (including trans-spliced) intron sequences, via a yet unknown mechanism (Cuenca *et al*. 2016, Grewe *et al*. 2016, Sloan *et al*. 2010). Interestingly, the introns loss is often followed by the presence of nucleotides in an edited state in the corresponding exonic loci. A proposed mechanism of plant mtDNA intron loss involves an RT-mediated model (Cuenca, *et al*. 2016, Grewe, *et al*. 2016, Sloan, *et al*. 2010). In canonical group II introns, homing occurs by a target DNA-primed RT mechanism initiated by an RNP complex containing the splicing factor and its excised intron (Cousineau *et al*. 1998). The ability of model group II introns to act as mobile RNAs (see *e*.*g*. (Lambowitz and Zimmerly 2011) (Schmitz-Linneweber, *et al*. 2015) is therefore of great mechanistic and evolutionary interest. The observation of reverse-transcription activity in mitochondria isolated from potato (*Solanum tuberosum*) tuber (Moenne *et al*. 1996), may support this intriguing possibility.

## Discussion

Mitochondria serve as principal sites for cellular energy metabolism, and also play key roles in the biosynthesis of numerous essential metabolites for the plant cell. The mitogenomes of land plants are notably larger and more complex in structure than their corresponding organelles in animals. Likewise, the expression of the mtDNA in plants is very complicated. Much of this control seem to occurs at the post-transcriptional level. RNA processing events that contribute to mtDNA expression in angiosperms, include RNA editing and the excision of numerous group II-type intron sequences, which disrupt the coding regions of many genes required in organellar translation and respiratory-mediated functions (Zmudjak and Ostersetzer-Biran 2017). Such a complexity may arise throughout the terrestrialization of plants, as a mean to control embryonic mitochondrial functions in order to optimize seed germination – a critical adaptive trait for life on land (Best, *et al*. 2020).

Transcriptomic data provide with comprehensive views into gene expression patterns and transcript’s levels (Wenz *et al*. 2009). Sequencing of yeast and human mitochondria transcriptomes provided with important insights into the importance of post-transcriptional processing for mitochondrial gene expression in eukaryotic cells (Mercer, *et al*. 2011, Turk, *et al*. 2013). Despite extensive investigations, the full landscapes of angiosperm’s mitochondrial transcriptomes, prerequisite for the elucidation of the basic plant mitochondria RNA biology, have not been developed yet.

Potential mitochondrial promoters, *e*.*g*. (Holec *et al*. 2006, Kühn *et al*. 2007, Kühn *et al*. 2009), transcript ends (Forner, *et al*. 2007, Ruwe, *et al*. 2016) and the processing of mtRNA have been previously reported. However, the molecular components of RNA processing and stability, complexes still remain to be elucidated. In this study, we analyzed the transcriptional landscapes of two key *Cruciferae* species, *Arabidopsis thaliana* (var. Col-0) and *Brassica oleracea* (var. botrytis), both of which are of vast importance for basic and organellar plant research. Relating mtRNA-seq and RT-qPCR data to the mitogenome database allowed us to elucidate transcript boundaries, analyze exon and intron sequences, RNA-editing events and ‘novel’ organellar transcripts, and access the relative abundances of mitochondrial transcripts. We provide here with details into the expression patterns and the identity of the 5’ and 3’ boundaries of the primary mitochondrial transcripts of Arabidopsis and cauliflower plants. The integrity of the data was further supported by RT-PCR and cDNA sequencing analyses. We further analyzed the accumulation of organellar transcripts encoding subunits of the respiratory complexes and genes found on polycistronic units. The mtRNA profiles indicate to differential expression of many OXPHOS genes, including of mRNA encoding to a specific protein complex. We therefore assume that the expression of respiratory subunits is likely to be regulated also at the translational or post-translational levels. In accordance with this assumption, ribosomal profiling experiments indicated to different abundances of mtRNAs in *Arabidopsis thaliana* (Planchard, *et al*. 2018), whereas analysis of the protein profiles provided with evidences for differential protein turnover rates in Arabidopsis mitochondria (Huang, *et al*. 2020). The unique properties of plant mitoribosomes (Waltz, *et al*. 2019) may associate with the differential expression of mitochondrial mRNAs seen in the two Brassicales species.

The studied transcriptome landscapes of Arabidopsis and cauliflower mitochondria provide with important aspects into mtRNA biology in angiosperm’s mitochondria, and we expect that these would serve as a valuable resource for the plant organellar community. Together, the revised mtRNA landscapes and previous analyses of the mitoribosome profiling experiments, proteomic studies, and the analysis of Arabidopsis mutants, provide with important insights into how all of the different parts of the mtDNA expression systems coordinate to provide the energy balance that is crucial to plant life.

## Acknowledgements

The authors confirm that they have no conflict of interest to declare. We would like to thank Mr. Roei Matan for his assistance with PCR and RT-PCR analyses. This work was supported by grants to O.O.B from the ‘Israeli Science Foundation’ ISF grants no. 741/15 and 1834/20.

